## Supplemental figures and tables for "Modulating the activities of chloroplasts and mitochondria promotes ATP production and plant growth"

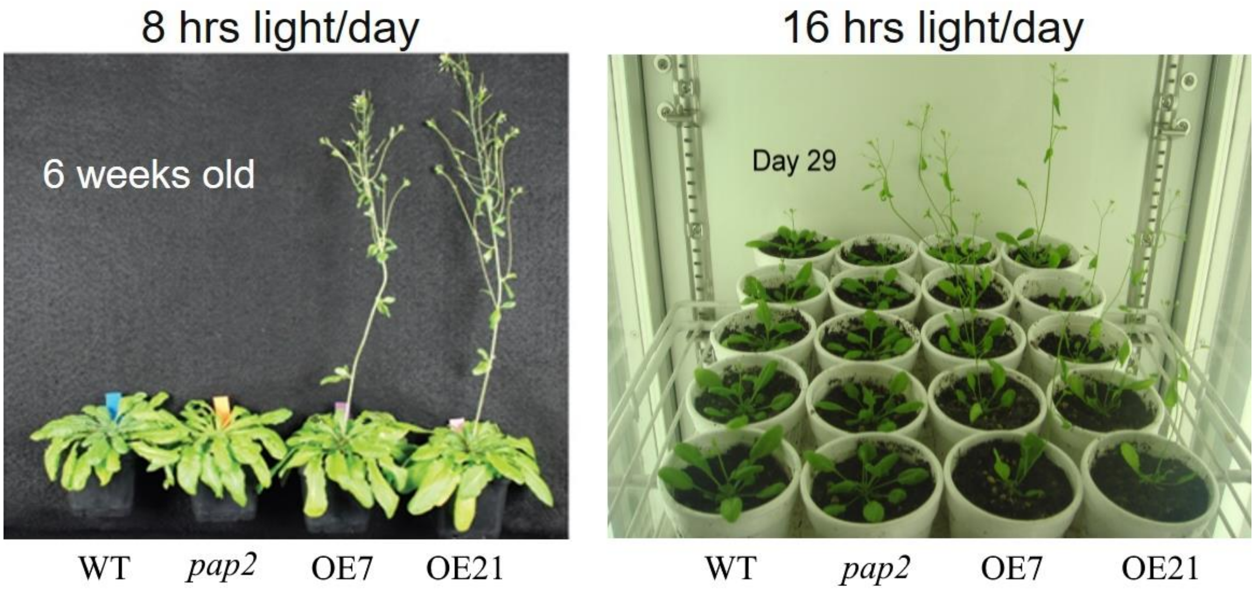


**Supplementary Figure S1. The OE lines grew faster under both SD (8h/16h) and LD (16h/8h) conditions.**

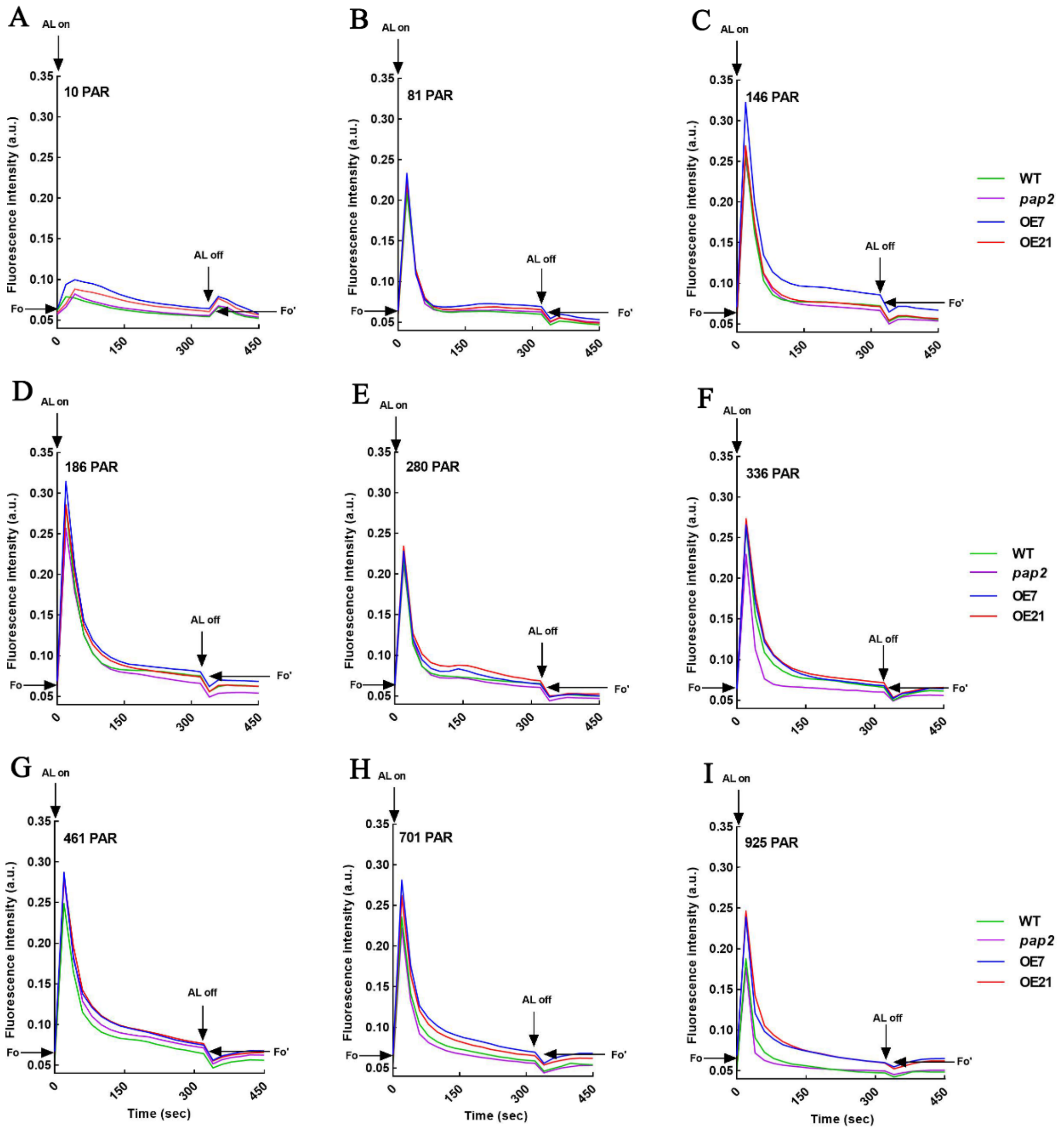


**Supplementary Figure S2. Fluorescence kinetics of WT, *pap2*, OE7 and OE21 lines.** 3-week-old plants were dark-acclimated for at least 1 hour before the measurement. Actinic light (AL) of 10, 81, 146, 186, 280, 336, 461, 701, 925 PAR was applied for 315 sec. Each data represents the mean of six biological replicates. **AL on**, actinic light on; **AL off**, actinic light off; **PAR**, photosynthetic active radiation (μmol photons $m^{-2}s^{-1}$); **Fo**, minimal fluorescence yield in the dark; **Fo'**, minimal fluorescence yield after illumination.

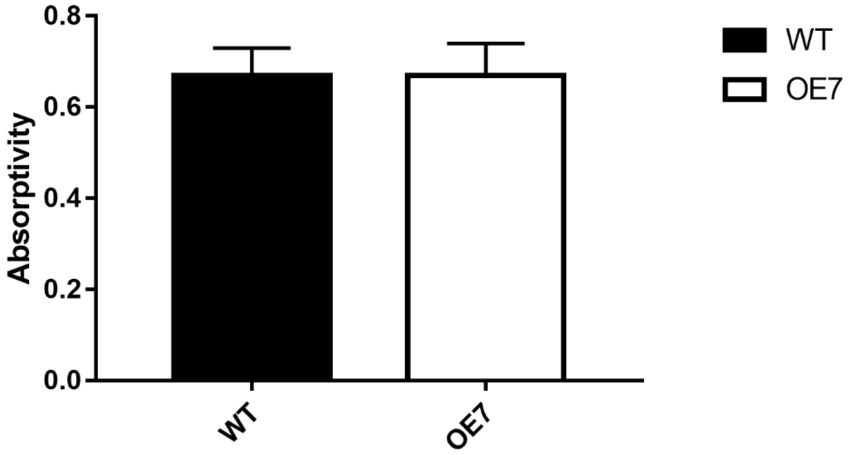


**Supplementary Figure S3. Leaf absorptivity is similar between WT line and AtPAP2 overexpression line (OE7).** Absorptivity of 20-day-old leaves were determined by IMAGING-PAM M-Series Maxi Version (WALZ, Germany). First an NIR (near-infrared)-remission image and then a R (red)-remission image was measured, the absorptivity was automatically calculated pixel by pixel according to the equation absorptivity = 1 - R/NIR. This approach is based on the empirical fact that pigments that contribute to the absorption of PAR (photosynthetic active radiation) do not show significant absorption bands in the NIR spectral region. On the other hands, pigments that absorb NIR are likely to also absorb Red light. Bar graph represents the mean value of absorptivity with SD (n = 63).

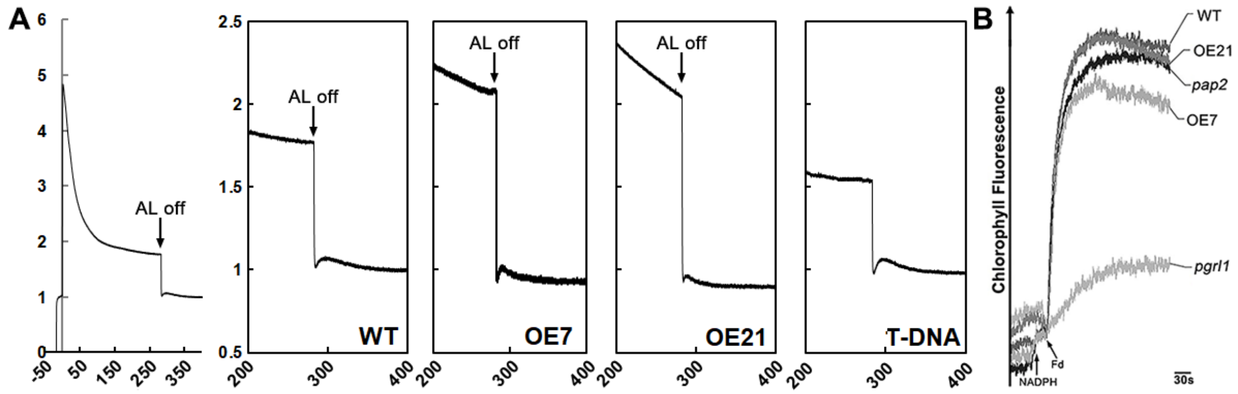


**Supplementary Figure S4.** ***In vivo*** **analysis of the cyclic electron flow rate of 20-day-old plants.** **A.** Post-illumination chlorophyll fluorescence transient for the measuring of NAD(P)H dehydrogenase (NDH)-CEF. Typical induction kinetics of chlorophyll fluorescence in a dark-acclimated leaf of WT Arabidopsis under illumination with actinic light (AL, 50 µmol photon $m^{-2}$ $s^{-1}$) were shown. Changes in chlorophyll fluorescence after the cessation of the actinic illumination were compared at a higher time resolution. Chlorophyll fluorescence was normalized to the original level in the dark ($F_0$). **B.** Ruptured chloroplast assay for the quantification of antimycin A-sensitive Fd-dependent CEF. After the addition of NADPH and Fd to freshly ruptured chloroplasts, the increase in chlorophyll fluorescence was measured (n = 3). The *pgrl1* mutant served as a negative control.

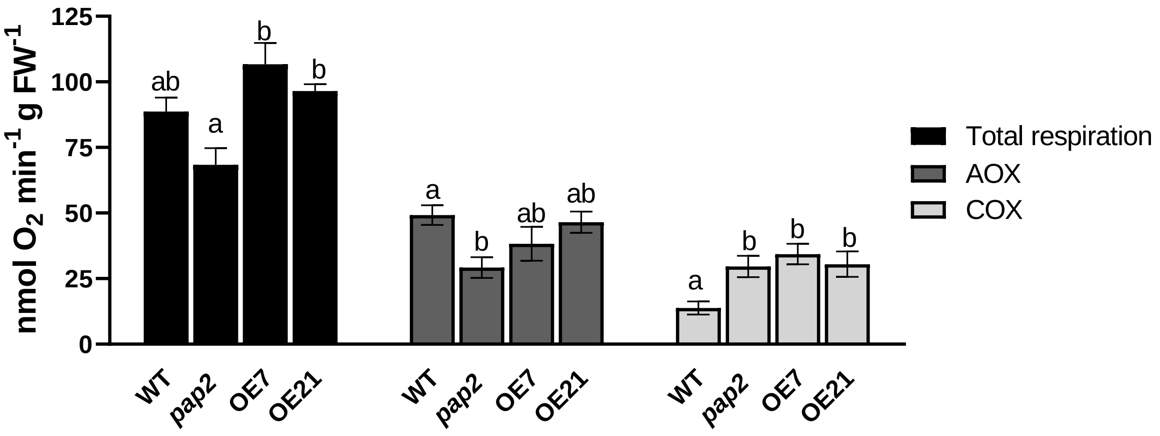


**Supplementary Figure S5. Respiratory rate of 21-day-old leaves.** Total respiration rate, AOX pathway respiration rate (the sensitivity of $O_2$ uptake to 10mM SHAM) and COX pathway respiration rate (the sensitivity of $O_2$ uptake to 4 mM KCN in the presence of 10mM SHAM) were expressed in nmol $O_2$ min$^{-1}$ g FW$^{-1}$ of WT, *pap2*, OE7 and OE21. Different letters indicate significant statistical differences analyzed by post-hoc Tukey's HSD test ($P < 0.05$) ($n = 5$; means ± SEM). **AOX**, alternative oxidase; **COX**, cytochrome c oxidase; **SHAM**, salicylhydroxamic acid; **KCN**, potassium cyanide.

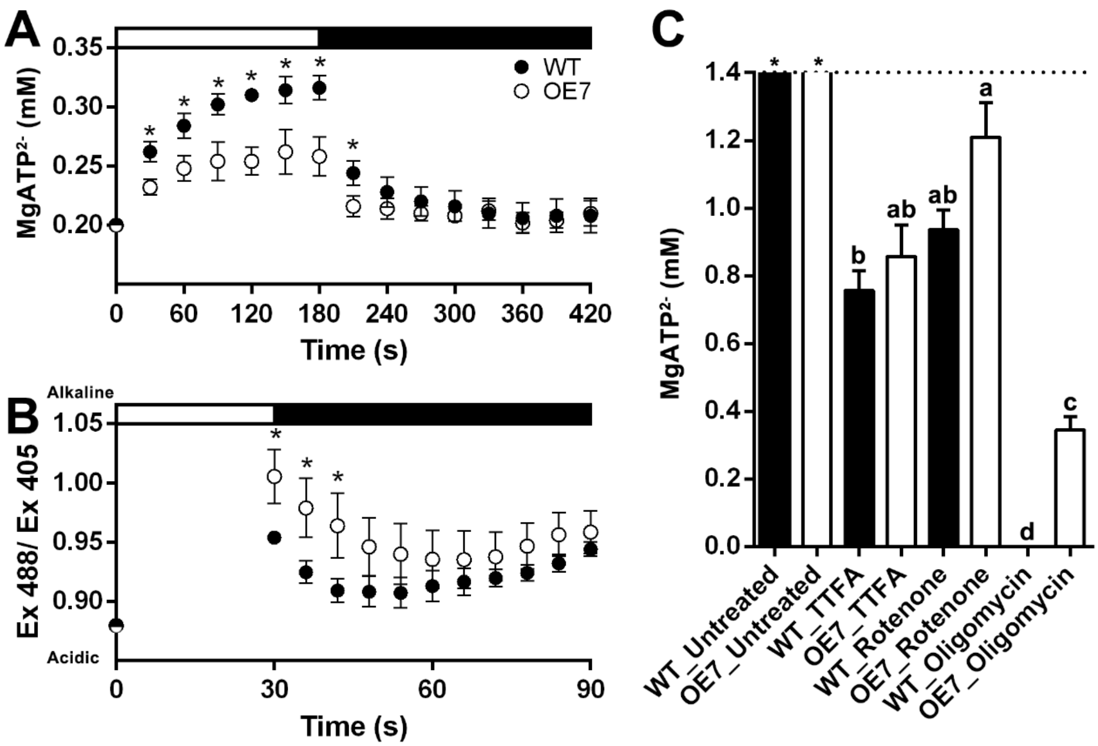


**Supplementary Figure S6. Assessing organelle activities using fluorescent sensors.** (a) Change in apparent $MgATP^{2-}$ concentration in the chloroplast stroma. 10-day-old WT and OE7 cotyledon expressing TKTP-AT1.03 were illuminated (296 µmol photon $m^{-2}$ $s^{-1}$) for 3 mins, followed by 5 mins in the dark. (b) Changes in the mitochondrial matrix pH sensor ratio (Ex 488/ Ex 405). 10-day-old WT and OE7 cotyledon expressing matrix-localized pH sensor, mt-cpYFP were illuminated (296 µmol photon $m^{-2}$ $s^{-1}$) for 30 s, followed by 1 min in the dark. Black and white bars indicate the dark and illumination periods, respectively. Average value of 6 independent measurements and SE are shown. The asterisks indicate significant differences between the WT and the OE line by one-way ANOVA with post-hoc Tukey HSD test ($p < 0.05$). Graphs were normalized to the initial point. (c) Effects of the inhibitors on cytosolic ATP concentration. Inhibitors of complex I (50 µM rotenone), complex II (100 µM TTFA) and complex V (10µM oligomycin) were applied to 10-day-old WT and OE7 seedling expressing C-AT1.03 seedlings by vacuum infiltration for 5 min. Before the measurements, the seedlings were incubated in the dark for 1 hour. Average value of 3 replicates is presented. Error bars are the standard error of the average $MgATP^{2-}$ concentration. Groups with significant difference by one-way ANOVA with post-hoc Tukey HSD test ($p < 0.05$) are indicated by different letters. The dashed line indicates the maximal $MgATP^{2-}$ concentration (1.4 mM) that can be reported by AT1.03 (Voon et al., 2018). The groups marked by asterisks are significantly different from the groups marked by letters but whether they have intergroup differences are not known.

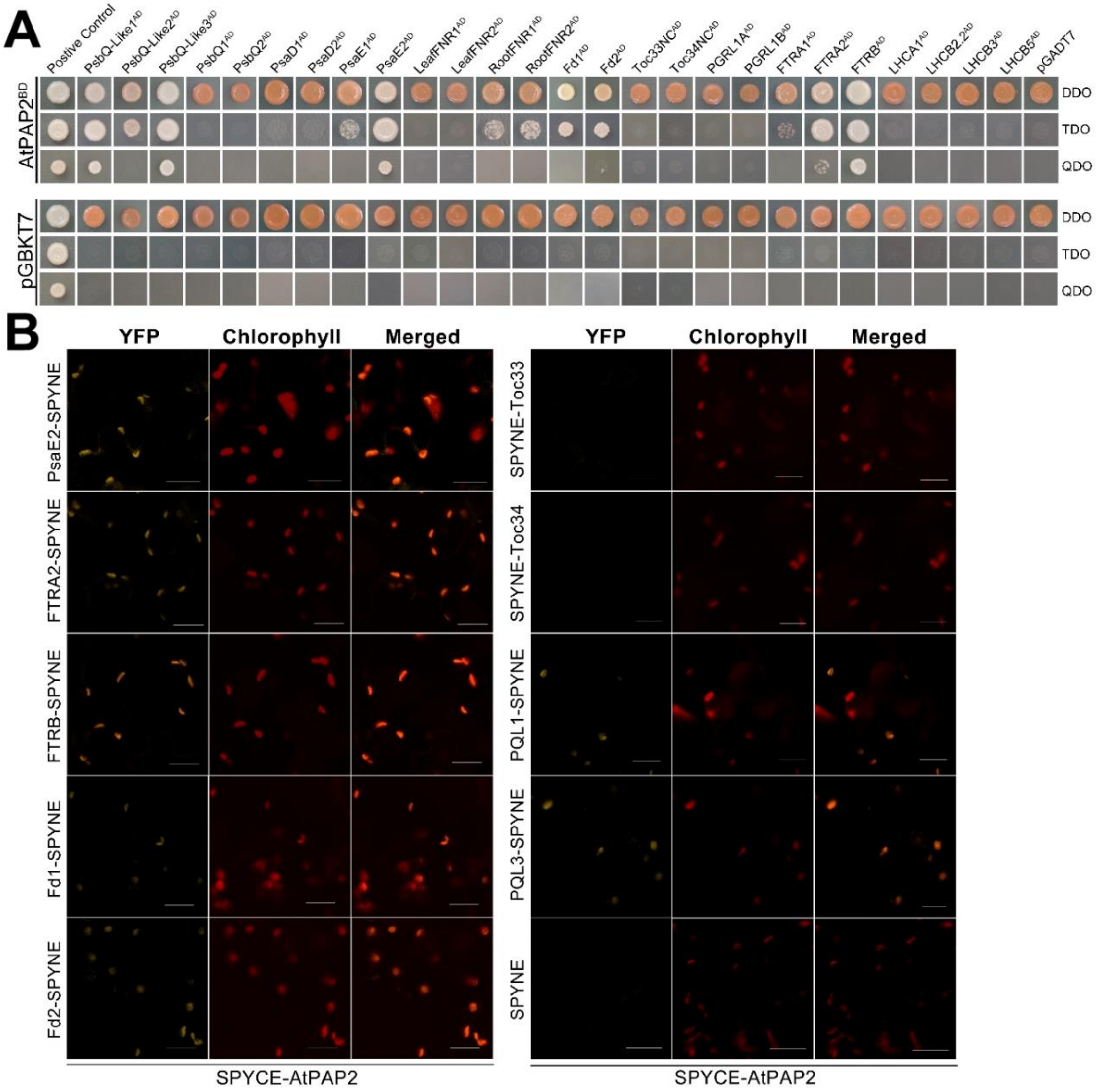


**Supplementary Figure S7. AtPAP2 selectively interacts with certain photosystem proteins.** **A**. Mature AtPAP2 protein without the signal peptide and C-terminus (a.a. 25–613, P2NC) interacted with several photosystem proteins (PQL1-AD, PQL2-AD, PQL3-AD, PsaE2-AD, FTRA2-AD and FTRB-AD). Blank vectors of BD (pGBKT7) and AD (pGADT7) were co-transformed with bait and prey, respectively, to assess auto-activation. The assays were carried on dropout medium of DDO (SD/-Leu/-Trp), TDO (SD/-Leu/-Trp/-His) and QDO (SD/-Ade/-His/-Leu/-Trp). **B**. BiFC assay of the interaction between AtPAP2 and candidate proteins. $YFP^N$-fused photosystem proteins (PsaE2, FTRA2, FTRB, Fd1, Fd2, PQL1, PQL3, Toc33 and Toc34) and $YFP^C$-AtPAP2 were transiently expressed in tobacco leaves. Empty SPYNE vectors were used as controls. Reconstituted YFP fluorescence was monitored at 514 nm (left panel with PMT detector emission bandwidth of 488–550 nm). Chlorophyll autofluorescence was excited at 458 nm (middle panel with PMT detector emission bandwidth of 650–710 nm). An overlay of the YFP signal and the chlorophyll autofluorescence is shown in the right panel. Scale bars: 20 µm. All images were captured using the same gain settings as the corresponding PMT channels. The representative images of 3 biological replicates are presented.

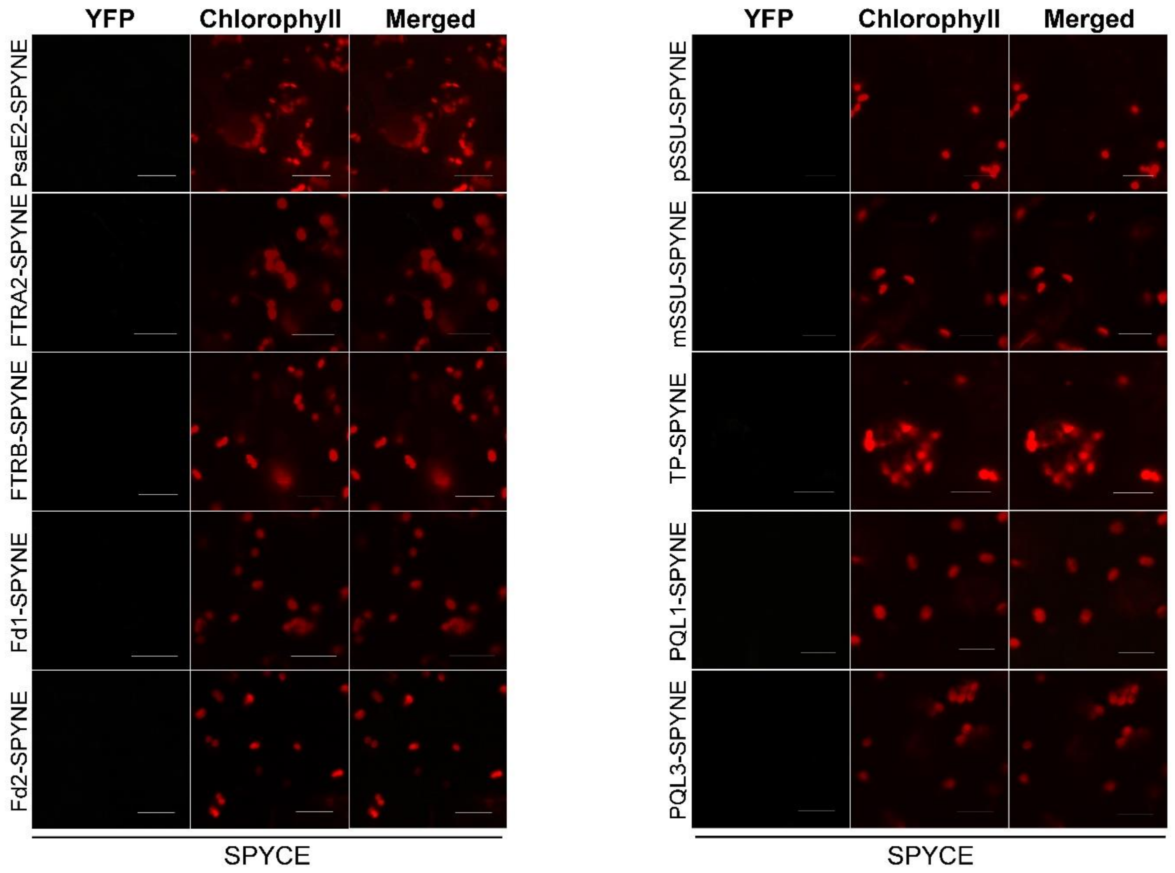


**Supplementary Figure S8. Results of BiFC negative controls.** YFPN-fused photosynthetic apparatus subunits (PsaE2, FTRA2, FTRB, PQL1, PQL3, Fd1, Fd2, SP, mSSU and pSSU) and empty SPYCE vector were transiently expressed in tobacco leaves as controls. Reconstituted YFP fluorescence was monitored at 514 nm (left panel with PMT detector emission bandwidth of 488–550 nm). Chlorophyll autofluorescence was excited at 458 nm (middle panel with PMT detector emission bandwidth of 650–710 nm). An overlay of the YFP signal and the chlorophyll autofluorescence is shown in the right panel. Scale bars: 20 μm. All of the images were captured at the same gain settings as the corresponding PMT channels.

**Supplementary Table S1. Photosynthetic pigment content.**

| | WT | *pap2* | OE7 | OE21 |
|---|---|---|---|---|
| Chlorophyll a | 8.43 + 0.44 [a] | 10.62 + 0.18 [b] | 6.53 + 1.12 [c] | 5.55 + 0.29 [c] |
| Chlorophyll b | 3.35 + 0.21 [a] | 3.96 + 0.87 [b] | 2.81 + 0.36 [ac] | 2.16 + 0.16 [c] |
| β−Carotene | 0.28 + 0.02 [a] | 0.30 + 0.07 [b] | 0.22 + 0.03 [c] | 0.18 + 0.00 [ac] |
| Lutein | 1.09 + 0.06 [a] | 1.28 + 0.07 [b] | 0.83 + 0.15 [c] | 0.68 + 0.02 [c] |
| Violaxanthin | 0.28 + 0.02 [a] | 0.29 + 0.08 [a] | 0.26 + 0.07 [a] | 0.18 + 0.01 [a] |

Pigments were extracted 8 hours after the light period from the leaves of 20-day-old Arabidopsis under LD regime. Values (mg $g^{-1}$ DW of tissues) marked by different letters are significantly different ($p < 0.05$) in the same row by one-way ANOVA analysis followed by HSD test. DW, dry weight.

**Supplementary Table S2. Chloroplast proteins identified in 2D BN PAGE.**

| Spot ID | Peptides (95%) | Unused peptide | Protein Name | Accession | Function | Average Ratio (OE/WT) |
|---|---|---|---|---|---|---|
| 1 | 103 | 162.86 | PsaA | ATCG00350 | PS I core protein | 1.00 |
| 1 | 80 | 98.48 | PsaB | ATCG00340 | PS I core protein | 1.00 |
| 2 | 38 | 69.11 | PsaA | ATCG00350 | PS I core protein | 0.82 |
| 2 | 38 | 62.91 | PsaB | ATCG00340 | PS I core protein | 0.82 |
| 23 | 72 | 78.56 | PsaD1 | AT4G02770 | PS I core protein | 0.97 |
| 12 | 23 | 29.77 | PsbA, D1 | ATCG00020 | PS II core protein | 0.78* |
| 6 | 95 | 82.02 | PsbB, CP47 | ATCG00680 | PS II core protein | 0.63** |
| 7 | 58 | 96.16 | PsbB, CP47 | ATCG00680 | PS II core protein | 0.72* |
| 8 | 51 | 64.74 | PsbC, CP43 | ATCG00680 | PS II core protein | 0.65** |
| 9 | 181 | 249.7 | PsbC, CP43 | ATCG00280 | PS II core protein | 0.80 |
| 12 | 41 | 58.68 | PsbD, D2 | ATCG00270 | PS II core protein | 0.78* |
| 13 | 108 | 136.85 | PsbO1, OE33 | AT5G66570 | PS II core protein | 6.83** |
| 14 | 59 | 83.54 | PsbO1, OE33 | AT5G66570 | PS II core protein | 8.64** |
| 15 | 15 | 26.96 | PsbO1, OE33 | AT5G66570 | PS II core protein | 3.02** |
| 21 | 22 | 25.47 | PsbS, NPQ4 | AT1G44575 | PS II core protein | 1.06 |
| 22 | 25 | 39.68 | PsbS, NPQ4 | AT1G44575 | PS II core protein | 1.01 |
| 16 | 145 | 173.1 | Lhcb1.4 | AT2G34430 | LHC II protein | 0.88 |
| 16 | 57 | 27.88 | Lhcb2.2 | AT2G05070 | LHC II protein | 0.88 |
| 17 | 91 | 140.01 | Lhcb4.1, CP29 | AT5G01530 | LHC II protein | 1.44 |
| 18 | 59 | 63.18 | Lhcb4.1, CP29 | AT5G01530 | LHC II protein | 0.79 |
| 18 | 33 | 33.7 | Lhcb1.4 | AT2G34430 | LHC II protein | 0.79 |
| 19 | 92 | 10.33 | Lhcb1.1 | AT1G29920 | LHC II protein | 0.92 |
| 19 | 93 | 129.87 | Lhcb1.4 | AT2G34430 | LHC II protein | 0.92 |
| 19 | 47 | 33.61 | Lhcb2.2 | AT2G05070 | LHC II protein | 0.92 |
| 19 | 37 | 52.97 | Lhcb5, CP26 | AT4G10340 | LHC II protein | 0.92 |
| 20 | 129 | 12.78 | Lhcb1.3 | AT1G29930 | LHC II protein | 0.84 |
| 20 | 125 | 190.01 | Lhcb1.4 | AT2G34430 | LHC II protein | 0.84 |
| 20 | 57 | 47.46 | Lhcb2.2 | AT2G05070 | LHC II protein | 0.84 |
| 20 | 56 | 92.39 | Lhcb5, CP26 | AT4G10340 | LHC II protein | 0.84 |
| 21 | 29 | 35.92 | Lhcb6, CP24 | AT1G15820 | LHC II protein | 1.06 |
| 22 | 71 | 99.62 | Lhcb6, CP24 | AT1G15820 | LHC II protein | 1.01 |
| 24 | 32 | 45.98 | PetC | AT4G03280 | Cytochrome B6f complex | 1.1 |
| 25 | 3 | 3.36 | PetD | ATCG00730 | Cytochrome B6f complex | 0.25** |
| 4 | 306 | 320.72 | atpA | ATCG00120 | ATP synthase | 0.32** |
| 5 | 325 | 381.21 | atpB | ATCG00480 | ATP synthase | 0.36** |
| 10 | 64 | 87.68 | atpC1 | AT4G04640 | ATP synthase | 0.60** |
| 11 | 58 | 62.97 | atpC1 | AT4G04640 | ATP synthase | 0.66** |
| 3 | 271 | 405.15 | RBCL | ATCG00490 | Calvin cycle | 4.07** |

Isolated chloroplast proteins from WT and OE7 were resolved by 2D BN/SDS-PAGE and the protein spots were extracted for MS/MS analysis. Protein candidates with the highest unused score and at least 2 unique peptides (95%) were shown. Average ratio is the mean of the value (OE/WT) from three biological replicates. Spots with fold change of ±1.2 or ±1.5 folds and $p < 0.05$ by one-way ANOVA are denoted by * and **, respectively.

**Supplementary Table S3. Summary of identified mitochondrial protein spots.**

| Spot ID | Peptides (95%) | Unused peptide | Protein name | Accession | Protein complex | Ratio (OE/WT) |
|---|---|---|---|---|---|---|
| 1 | 327 | 328.85 | Heat Shock Protein 60-3B | AT3G23990.1 | HSP60 | 1.07 |
| 1 | 262 | 158.77 | Heat Shock Protein 60-2 | AT2G33210.1 | HSP60 | 1.07 |
| 2 | 244 | 288.80 | MPPbeta | AT3G02090.1 | Complex III | 0.99 |
| 3 | 295 | 358.38 | ATP synthase beta-subunit | AT5G08690.1 | Complex V | 0.52** |
| 3 | 292 | 4.48 | ATP synthase beta-subunit | AT5G08680.1 | Complex V | 0.52** |
| 3 | 239 | 312.37 | ATP synthase alpha-subunit | AT2G07698.1 | Complex V | 0.52** |
| 4 | 217 | 287.35 | ATP synthase beta-subunit | AT5G08690.1 | Complex V | 0.43** |
| 4 | 209 | 6.00 | ATP synthase beta-subunit | AT5G08680.1 | Complex V | 0.43** |
| 4 | 8 | 234.30 | ATP synthase alpha-subunit | AT2G07698.1 | Complex V | 0.43** |
| 5 | 622 | 637.08 | RBCL | ATCG00490.1 | | 2.33** |
| 6 | 342 | 349.23 | ATP synthase beta-subunit | AT5G08690.1 | Complex V | 0.56** |
| 6 | 335 | 6.00 | ATP synthase beta-subunit | AT5G08680.1 | Complex V | 0.56** |
| 6 | 204 | 220.55 | ATP synthase alpha-subunit | AT2G07698.1 | Complex V | 0.56** |
| 7 | 245 | 315.84 | Insulinase protein | AT1G51980.1 | Complex III | 1.08 |
| 7 | 73 | 11.84 | MPPalpha | AT3G16480.1 | Complex III | 1.08 |
| 7 | 14 | 22.21 | MPPbeta | AT3G02090.1 | Complex III | 1.08 |
| 8 | 242 | 335.95 | ATP synthase gamma-subunit (ATP3) | AT2G33040.1 | Complex V | 0.64** |
| 8 | 25 | 41.00 | ATP Synthase subunit 1 (ATP1) | ATMG01190.1 | Complex V | 0.64** |
| 8 | 20 | 35.36 | ATP synthase beta-subunit | AT5G08690.1 | Complex V | 0.64** |
| 9 | 19 | 28.61 | ATP synthase alpha-subunit | AT2G07698.1 | Complex V | 0.65** |
| 9 | 32 | 56.70 | ATP synthase gamma-subunit (ATP3) | AT2G33040.1 | Complex V | 0.65** |
| 9 | 29 | 50.43 | ATP synthase beta-subunit | AT5G08690.1 | Complex V | 0.65** |
| 10 | 87 | 131.85 | ATP synthase gamma-subunit (ATP3) | AT2G33040.1 | Complex V | 0.64** |
| 10 | 49 | 72.64 | ATP synthase beta-subunit | AT5G08670.1 | Complex V | 0.64** |

Isolated mitochondrial proteins from WT and OE7 were resolved by 2D BN/SDS-PAGE and the mitochondrial protein spots were extracted for MS/MS analysis. Protein candidates with the highest unused score and at least 2 unique peptides (95%) were shown. Average ratio is the mean of the value (OE/WT) from three biological replicates. Spots with fold change of ±1.2 or ±1.5 folds and $p < 0.05$ by one-way ANOVA are denoted by * and **, respectively.

**Supplementary Table S4. Respiration rates of different complexes of WT and OE7 mitochondria.**

| Enzyme | Substrate | ADP | Rotenone | WT | OE7 | Change |
|---|---|---|---|---|---|---|
| Complex I | deamino-NADH | - | - | 30 ± 6 | 33 ± 5 | ns |
| | | - | + | 8 ± 3 | 8 ± 5 | ns |
| Complex II | Succinate | - | - | 49 ± 3 | 48 ± 23 | ns |
| | | + | - | 75 ± 4 | 92 ± 10 * | 1.23X |
| NdEx | NADH | - | - | 45 ± 4 | 45 ± 12 | ns |
| | | + | - | 65 ± 4 | 68 ± 4 | ns |
| NdIn | Malate + Glutamate | - | + | 39 ± 4 | 46 ± 6 | ns |
| | | + | + | 64 ± 8 | 85 ± 4 * | 1.32X |

Oxygen consumption rates of complex I (rotenone-sensitive deamino-NADH oxidation), complex II, external NADH dehydrogenase, internal NADH dehydrogenase (rotenone-insensitive NADH oxidation) were expressed as nmol $min^{-1}$ $mg^{-1}$ protein.

* indicates statistically significant difference ($p < 0.05$, $n \geq 3$) and ns indicates not significant difference between WT and OE7 by Student's t test.

**Supplementary Table S5. AtPAP2-interacting proteins identified by Y2H library screening.**

| AGI code | No. of clones | a.a. residues | Sequence in clone | Phosphorylation sites (1-50) | pI | Proteins | Experimental location |
|---|---|---|---|---|---|---|---|
| At1g14150 | 5 | 190 | All are full length | T9,Y13,T46 | 8.93 | PsbQ-like (PQL1) | Plastid |
| At1g18640 | 5 | 695 | All are full length | *T6* | 6.29 | 3-Phosphoserine phosphatase | Plastid |
| At1g26220 | 1 | 197 | Full length | T9,T27,S31,Y46,S47,S49 | 9.79 | Acyl-CoA N-acyltransferases superfamily protein | Plastid |
| At1g29900 | 2 | 1187 | aa.771-End | *S18,S19,S26,S31,S33* | 5.45 | Carbamoyl phosphate synthetase B (CARB) | Plastid |
| At1g30510 | 1 | 382 | aa.135-End | S15,T49 | 8.85 | Ferredoxin--NADP+ reductase (RFNR2) | Plastid |
| At2g04700 | 5 | 146 | All are full length | S15,T20,T34,S48 | 7.98 | Ferredoxin-thioredoxin reductase B | Plastid |
| At2g05990 | 1 | 390 | aa.149-End | S17,S19,S20,Y30 | 9.36 | Enoyl-ACP reductase | Plastid |
| At2g20260 | 1 | 145 | Full length | S24,S25 | 10.54 | Subunit E of photosystem I (PsaE2) | Plastid |
| At2g22360 | 1 | 442 | aa.228-End | Y25,S32 | 9.49 | DNAJ heat shock family protein A6 | Plastid |
| At2g34860 | 1 | 186 | Full length | S16,S39,S42,S45,S47,S49 | 9.25 | DnaJ/Hsp40 cysteine-rich domain protein | Plastid |
| At2g35500 | 1 | 387 | Full length | *S14,S34,S35* | 6.19 | Shikimate kinase-like 2 (SKL2) | Plastid |
| At4g20760 | 1 | 298 | Full length | S14,S34,S37,T32 | 10.24 | NAD(P)-binding Rossmann-fold protein | Plastid |
| At4g21860 | 1 | 202 | Full length | T7,S34,S45,S46 | 8.68 | Methionine sulfoxide reductase B2 | Plastid |
| At4g25370 | 1 | 238 | Full length | S26,S28,S29,T36 | 9.78 | Double Clp-N motif protein | Plastid |
| At4g35860 | 1 | 211 | Full length | T49 | 7.03 | GTP-binding protein GB2 (GB2) | Plastid |
| At5g06340 | 3 | 227 | All are full length | S37,S50 | 8.77 | Nudix hydrolase 27 (NUDX27) | Plastid |
| At5g08280 | 1 | 382 | Full length | *T26,S44* | 8.62 | Hydroxymethylbilane synthase | Plastid |
| At5g11450 | 4 | 297 | All are full length | S20,Y27,T43,S45,S47 | 9.06 | PPD5 | Plastid |

| | | | | | | | |
|---|---|---|---|---|---|---|---|
| At5g11650 | 1 | 390 | aa.323-End | *T12* | 8.73 | Alpha/beta fold hydrolase family protein | Plastid |
| At5g19370 | 4 | 299 | All are full length | T7,S9,S13,T30 | 8.62 | Rhodanese-like Peptidyl-prolyl cis-trans isomerase | Plastid |
| At5g21222 | 1 | 831 | aa.523-End | Y13,T33,T35 | 8.01 | Serine/threonine protein kinase | Plastid |
| At5g38430 | 2 | 181 | Full length | S14,S32,T36 | 7.83 | Ribulose bisphosphate carboxylase small chain 1B | Plastid |
| At5g66120 | 2 | 442 | aa.1-344 and 1-343 | T14,T31,Y50 | 7.53 | 3-dehydroquinate synthase | Plastid |
| At4g33520 | 1 | 949 | Full length | T14 | 9.02 | Copper-transporting ATPase PAA1 (PAA1) | Plastid inner membrane |
| At1g53670 | 1 | 202 | Full length | Y23,Y48 | 9.20 | Methionine sulfoxide reductase B1 | Plastid stroma |
| At2g20270 | 1 | 206 | Full length | T45 | 8.94 | Monothiol glutaredoxin-S12 | Plastid stroma |
| At2g38270 | 4 | 293 | All are full length | S15,T31,T37,S41 | 8.07 | Monothiol glutaredoxin 16 | Plastid stroma |
| At2g36460 | 2 | 358 | aa.76-End and aa.112-End | S32,T33,T35,S42 | 7.46 | Fructose-bisphosphate aldolase, class I | Plastid, cytosol, PM, extracellular |
| At1g18080 | 1 | 327 | aa.53-End | T19,T23 | 7.81 | RACK1A | Plastid, cytosol, PM, nucleus |
| At1g70580 | 1 | 481 | aa.145-End | Y8,Y27 | 6.52 | Alanine-2-oxoglutarate aminotransferase 2 | Plastid, Peroxisome |
| At5g16370 | 1 | 552 | aa. 871-1766 | S11,T15 | 6.97 | Acyl activating enzyme 5 (AAE5) | Plastid, Peroxisome |
| At4g25130 | 3 | 258 | All are full length | *S7* | 8.92 | Methionine sulfoxide reductase | Plastid, stroma |
| At2g35240 | 1 | 232 | Full length | S19,S26,S40,Y49 | 9.14 | pMORF6 | Mitochondria |
| At5g40770 | 1 | 277 | aa.47-End | T34 | 7.89 | Prohibitin 3 (PHB3) | Mitochondria |
| At4g35830 | 1 | 898 | aa.467-End | Y41 | 6.35 | Aconitate hydratase 1 (ACO1) | Mitochondria and cytosol |
| At1g05060 | 1 | 253 | Full length | Y16,S32,T35,S41,S42 | 10.02 | Uncharacterized protein | N/A |
| At1g29810 | 1 | 187 | Full length | T42,S47,T49,T50 | 10.37 | Transcriptional coactivator/pterin dehydratase | N/A |

| | | | | | | | |
|---|---|---|---|---|---|---|---|
| At1g31817 | 1 | 314 | aa.48-End | S31,S33,S37,S48 | 11.15 | 30S ribosomal protein S11 | N/A |
| At1g53140 | 1 | 817 | aa.620-End | Y7,T10,T14 | 7.47 | Dynamin related protein 5A (DRP5A) | N/A |
| At2g38080 | 1 | 558 | aa.172-534 | T49 | 9.60 | Laccase/Diphenol oxidase family protein | N/A |

Experimental location: determined by GFP or MS analysis (SUBA3, http://suba.plantenergy.uwa.edu.au/)
Phosphorylation sites (1-50): Experimental determined phophorylation sites in the first 50 a.a. residues were retrieved from Phosphat 4.0 (http://phosphat.uni-hohenheim.de/phosphat.html)
Predicted location: predicted by SUBA3 (http://suba.plantenergy.uwa.edu.au/)
Predicted phosphorylation sites (1-50) are shown in italics. The prediction was carried out at Phosphat 4.0 or NetPhos 2.0 (http://www.cbs.dtu.dk/services/NetPhos/)

**Supplementary Table S6. Primer list.**

| Vector (Purpose) | Primer Name (AGI code) | Primer Sequence (5'-3')[1] |
|---|---|---|
| pGBKT7 | 25-613aaAtPAP2 | F: TTCTCATATGACCATTTCAATTTCCCC |
| (Yeast two-hybrid) | (AT1G13900) | R: GCATGTCGACCAGCATTAGATTCTGATTTTC |
| pGADT7 | 42-181aaSSU1B | F: ATTCCATATGGACATTACTTCCATCACAAGC |
| (Yeast two-hybrid) | (AT5G38430) | R: ATTACTCGAGAGCATCAGTGAAGCTTGGGG |
| pGADT7 | 1-50aaSSU1B | F: ATTACATATGGCTTCCTCTATGCTCTCCT |
| (Yeast two-hybrid) | (AT5G38430) | R: ATTACTCGAGCCCATTGCTTGTGATGGAAG |
| pGADT7 | PsbQ1 | F: ATTAGAATTCATGGCTTCGATGGGTGGATT |
| (Yeast two-hybrid) | (AT4G21280) | R: CTAGCTCGAGTTAACCAAGCTTGGCAAGAACT |
| pGADT7 | PsbQ2 | F: TTAGCATATGATGGCTCAAGCAGTGACTTC |
| (Yeast two-hybrid) | (AT4G05180) | R: ATTAGAGCTCTTAACCGAGCTTGGCAAGAAC |
| pGADT7 | PsbQ-Like1 | F: ATTGCATATGATGAGCTCCTTCACCACCAC |
| (Yeast two-hybrid) | (AT1G14150) | R: ATCGCTCGAGTTAAGCAAGAAACTCCACAAC |
| pGADT7 | PsbQ-Like2 | F: TGTCCATATGATGGCTCACTTCATAGACCT |
| (Yeast two-hybrid) | (AT3G01440) | R: TAATCTCGAGTCATGCCATTCTGGTCATTAC |
| pGADT7 | PsbQ-Like3 | F: TAATCATATGATGGCGATTTCAAAGCCACC |
| (Yeast two-hybrid) | (AT2G01918) | R: ACCGCTCGAGTTAAATTCGAGGGAAGATATC |
| pGADT7 | PsaD1 | F: ATTACATATGATGGCAACTCAAGCCGCCGG |
| (Yeast two-hybrid) | (AT4G02770) | R: GCTACTCGAGTTACAAATCATAACTTTGTTTGCCAG |
| pGADT7 | PsaD2 | F: ATTACATATGATGGCAACTCAAGCCGCCGG |
| (Yeast two-hybrid) | (AT1G03130) | R: GTCTCGAGTTACAAATCATAAGATTGTTTCCCAGTG |
| pGADT7 | PsaE1 | F: ATTACATATGATGGCGATGACGACAGCATC |
| (Yeast two-hybrid) | (AT4G02770) | R: CTTACTCGAGTTAAGCTGCAACTTCTTCGACCT |
| pGADT7 | PsaE2 | F: ATTACATATGATGGCGATGACGTCAGCAGC |
| (Yeast two-hybrid) | (AT2G20260) | R: GTTACTCGAGTCATTTTACTTCTTCCACCTCGTCC |
| pGADT7 | LeafFNR1 | F: TTAGCATATGATGGCTGCTGCTATAAGTGC |
| (Yeast two-hybrid) | (AT5G66190) | R: TACGCTCGAGTTAGTAGACTTCAACATTCCACTG |
| pGADT7 | LeafFNR2 | F: ATTACATATGATGGCGACTACCATGAATGC |
| (Yeast two-hybrid) | (AT1G20020) | R: TCTACTCGAGTCAGTAGACTTCAACGTTCCATT |
| pGADT7 | RootFNR1 | F: ATTGCATATGATGGCTCTCTCAACTACTCC |
| (Yeast two-hybrid) | (AT3G05390) | R: GCTACTCGAGTCAATACACTTCAACATGCC |
| pGADT7 | RootFNR2 | F: ATTGCATATGATGTCTCACTCTGCTGTTTCT |
| (Yeast two-hybrid) | (AT1G30510) | R: CTTACTCGAGTCAATAGACTTCAACGTGCC |
| pGADT7 | LeafFd1 | F: ATTACATATGATGGCTTCCACTGCTCTCTCC |
| (Yeast two-hybrid) | (AT1G10960) | R: ATTACTCGAGTTACATAATGGCTTCTTCTTTGTGG |
| pGADT7 | LeafFd2 | F: ATTGCATATGATGGCTTCCACTGCTCTCTC |
| (Yeast two-hybrid) | (AT1G60950) | R: GTCACTCGAGTTAAACAATGTCTTCTTCTTTGTGG |
| pGADT7 | 1-251aaToc33 | F: ATTACATATGATGGGGTCTCTCGTTCGTG |
| (Yeast two-hybrid) | (AT1G02280) | R: ATTACCCGGGTTAGTCTACATGAATTGCTTTCCTC |
| pGADT7 | 1-254aaToc34 | F: ATTACATATGATGGCAGCTTTGCAAACGCT |
| (Yeast two-hybrid) | (AT5G05000) | R: ATTACCCGGGTTACTTGTCAACATGAATCGCCTT |
| pGADT7 | PGR5-Like1A | F: ATTGCATATGATGGGTAGCAAGATGTTGTT |
| (Yeast two-hybrid) | (AT4G22890) | R: TAATCTCGAGTTAAGCTTGGCTTCCTTCTG |
| pGADT7 | PGR5-Like1B | F: ATTGCATATGATGGCTTTTACTCTAACAATCC |
| (Yeast two-hybrid) | (AT4G11960) | R: AACTGAGCTCTTAAGCTTTCCCTCCTTCTG |
| pGADT7 | FTRA1 | F: ATTGCATATGATGAGTAGCCAAATCGCTTTGT |
| (Yeast two-hybrid) | (AT5G23440) | R: TACTGAGCTCTCACTGATCAATGAACTCGAACTC |
| pGADT7 | FTRA2 | F: ATTGCATATGATGACTAACAGTTACGCTCTGTC |
| (Yeast two-hybrid) | (AT5G08410) | R: ATTACTCGAGTCACGGATCAATTAACTCGAAC |
| pGADT7 | FTRB | F: AGCTGAATTCATGAATCTTCAAGCTGTTTC |
| (Yeast two-hybrid) | (AT2G04700) | R: ATCGCTCGAGTCACATGTTAGCTGTAGTTTC |

| | | |
|---|---|---|
| pGADT7 | LHCA1 | F: TAATCATATGGCGTCGAACTCGCTTAT |
| (Yeast two-hybrid) | (AT3G54890) | R: TAATCATATGTTAGTTGAAAGGGATAACAAT |
| pGADT7 | LHCB2.2 | F: TAATCATATGGCCACATCAGCTATCCAAC |
| (Yeast two-hybrid) | (AT2G05070) | R: TAATCTCGAGTTACTTTCCGGGGACAAAGTT |
| pGADT7 | LHCB3 | F: TAATCATATGGCATCAACATTCACGAGC |
| (Yeast two-hybrid) | (AT5G54270) | R: TAATCTCGAGTTAAGCTCCAGGTGCAAACT |
| pGADT7 | LHCB5 | F: TAATCATATGGCGTCTTTGGGTGTGTC |
| (Yeast two-hybrid) | (AT4G10340) | R: TAATCTCGAGTTAGAGAGTGGGAGCTCTCTC |
| SPYCE | AtPAP2 | F: ATCGACTAGTATGATCGTTAATTTCTCTTTCTTC |
| (BiFC) | (AT1G13900) | R: GAATACTAGTTTATGTCTCCTCGTTCTTGACTG |
| SPYCE | Multiple cloning site | F: ATTAGAGCTCGTTAACCGGGCTCAGGCCT |
| (BiFC) | | R: ATTAGAGCTCCCCGGGAGCGGTACCCTC |
| SPYCE | $YFP^C$ | F: TAACTCTAGAATGTACCCATACGATGTTCCAG |
| (BiFC) | | R: ATTAGAGCTCCTTGTACAGCTCGTCCATG |
| SPYNE | PsaE1 | F: GAACTCTAGAATGGCGATGACGACAGCAT |
| (BiFC) | (AT4G28750) | R: ATTACTCGAGAGCTGCAACTTCTTCGACCTC |
| SPYNE | PsaE2 | F: ATTATCTAGAATGGCGATGACGTCAGCAGC |
| (BiFC) | (AT2G20260) | R: TAATCTCGAGTTTTACTTCTTCCACCTCGTCCAAT |
| SPYNE | FTRA1 | F: ATTATCTAGAATGAGTAGCCAAATCGCTTT |
| (BiFC) | (AT5G23440) | R: ATTACTCGAGCTGATCAATGAACTCGAACTC |
| SPYNE | FTRA2 | F: CTAGTCTAGAATGACTAACAGTTACGCTCTGTC |
| (BiFC) | (AT5G08410) | R: ATTACTCGAGCGGATCAATTAACTCGAACTC |
| SPYNE | FTRB | F: ATTATCTAGAATGAATCTTCAAGCTGTTTCTTG |
| (BiFC) | (AT2G04700) | R: ATTACTCGAGCATGTTAGCTGTAGTTTCTTTTATT |
| SPYNE | LeafFd1 | F: ATTATCTAGAATGGCTTCCACTGCTCTCTCC |
| (BiFC) | (AT1G10960) | R: TAATCTCGAGCATAATGGCTTCTTCTTTGTGG |
| SPYNE | LeafFd2 | F: ATTGTCTAGAATGGCTTCCACTGCTCTCTC |
| (BiFC) | (AT1G60950) | R: ATTACTCGAGAACAATGTCTTCTTCTTTGTGGG |
| SPYNE | Toc33 | F: ATTATCTAGAATGGGGTCTCTCGTTCGTGAAT |
| (BiFC) | (AT1G02280) | R: CCTATCTAGAAAGTGGCTTTCCACTTGTCTTGAT |
| SPYNE | Toc34 | F: ATTATCTAGAATGGCAGCTTTGCAAACGCT |
| (BiFC) | (AT5G05000) | R: GAATTCTAGAAGACCTTCGACTTGCTAAACCG |
| SPYNE | PsbQ-Like1 | F: ATTGACTAGTATGAGCTCCTTCACCACCAC |
| (BiFC) | (AT1G14150) | R: ATTACTCGAGAGCAAGAAACTCCACAACATT |
| SPYNE | PsbQ-Like3 | F: ATTCACTAGTATGGCGATTTCAAAGCCACC |
| (BiFC) | (AT2G01918) | R: ATTACTCGAGAATTCGAGGGAAGATATCATCGAG |
| SPYNE | 42-181aaSSU1B | F: ATTCTCTAGAATGGACATTACTTCCATCACAAGC |
| (BiFC) | (AT5G38430) | R: ATTACTCGAGAGCATCAGTGAAGCTTGGGG |
| SPYNE | 1-50aaSSU1B | F: ATTATCTAGAATGGCTTCCTCTATGCTCTCCT |
| (BiFC) | (AT5G38430) | R: ATTACTCGAGCCCATTGCTTGTGATGGAAG |
| pCXSN | TOC33overlap | F: ATCAGTGGGAGCAAAGTAGATGGATCTTACTCT |
| (Transgenic line) | (At1g02280) | R: GTAAGATCCATCTACTTTGCTCCCACTGATTAC |

[1] Restricted sites are indicated by underline.

**Supplementary Methods**

**Determination of the chlorophyll content using an HPLC analysis.** Leaves (30-50 mg) were freeze-dried and extracted 3 times using 100% acetone (500 μl, 250 μl and 250 μl). The mixture was combined and centrifuged twice to eliminate the insoluble substances prior to the HPLC analysis. The pigment extract was separated and analysed (20 μL aliquots) on a Waters Spherisorb 5 μm ODS2 (4.6 × 250 mm) analytical column (Waters, USA) at room temperature. The fluids were eluted at a flow rate of 1.2 ml $min^{-1}$ with a linear gradient from 100% solvent A [acetonitrile/methanol/0.1 M TRIS–HCl (pH 8.0), 84:2:14, v/v/v] to 100% solvent B (methanol/ethyl acetate, 68:32, v/v) over a 15-min period, followed by 10 min of 100% solvent B. The tridimensional chromatogram was recorded from 250 to 700 nm. The chromatographic peaks were identified by comparing the retention times and spectra to known standards provided by Sigma (USA) and Wako (Japan). The pigments were finally quantified by integrating the peak areas and converting them to concentrations according to the comparisons. The amounts of lutein, β–carotene, violaxanthin, chlorophyll a (Ca) and chlorophyll b (Cb) were calculated as follows: lutein: $Y = 2.56 \times 10^8 X - 153665$, $R = 0.99577$, $p < 0.0001$; β-carotene: $Y = 6.57 \times 10^8 X – 61818$; violaxanthin: $Y = 2.68 \times 10^8 X + 236.6$; Chl a: $Y = 3.30 \times 10^7 X + 38832$; and Chl b: $Y = 7.31 \times 10^7 X + 29646$. Y = area, X = mg/ml.

**Transmission electron microscopy (TEM).** Plant materials (5th to 7th true leaves of 19-day-old plants at the beginning of the light period) were fixed with 2.5% (v/v) glutaraldehyde in 75 mM sodium cacodylate and 2 mM $MgCl_2$, pH 7.0, for 1 hour at 25 °C and post-fixed for 2 hours with 1% (w/v) osmium tetroxide in a fixative buffer at 25 °C. After two washing steps with distilled water, the pieces were dehydrated and then embedded in Spurr's low-viscosity resin. Ultrathin sections of 50–70 nm were cut with a diamond knife. The sections were post-stained with aqueous uranyl acetate/lead citrate. The TEM examination was performed under a Hitachi H-7650 transmission electron microscope with a charge-coupled device camera (Hitachi High-Technologies, Japan) operating at 80 kV.

**2D Blue Native PAGE DIGE.** Leaves from 20-day-old soil-grown WT and OE7 plants at the middle of the day (T = 8) were used as the starting materials. The chloroplast (Lamkemeyer et al., 2006) and mitochondria isolation (Lee et al., 2008) were performed as previously described. Cy

Dye labelling of the chloroplast or mitochondrial proteins was carried out as previously described with modifications (Heinemeyer et al., 2009). In total, 100 µg chloroplast/mitochondrial proteins were centrifuged for 10 min at 4 °C at 1000 g using an Eppendorf centrifuge. The sedimented chloroplasts or mitochondria were re-suspended in a 10 µl solubilization buffer containing 30 mM HEPES, pH 7.4, 150 mM potassium acetate, 10% [v/v] glycerol, supplemented with 2% [w/v] beta-dodecyl maltoside (chloroplasts) or 5% [w/v] digitonin (mitochondria). The solubilized proteins were centrifuged for 20 min at 4 °C at full speed to remove the insoluble material. The supernatant containing the solubilized protein complexes was supplemented with 10 µl of the solubilization buffer containing 30 mM HEPES, pH 10, 150 mM potassium acetate, 10% [v/v] glycerol to adjust the pH value of the protein solution to approximately 8.5, which is a prerequisite for efficient labelling. Cy2, Cy3 and Cy5 NHS ester minimal dyes (Lumiprobe Corporation, USA) were reconstituted into 400 pmol/µl in DMF according to the manufacturer's instructions, and 2 µl of the Cy dye were added to the solubilized chloroplast or mitochondrial proteins. The WT was labelled using a Cy3 dye, and the OE was labelled with the Cy5 dye, while the internal standard (consisting of a pooled sample comprising an equal amount of all samples in the experiment) was labelled with Cy2. The labelling occurred for 30 min in the dark on ice and was stopped by adding 1 µl of a Lysine solution (10 mM). Finally, the sample was supplemented with 3 µl 5% (w/v) Serva Blue G (750 mM aminocaproic acid, 5% [w/v] Coomassie 250 G) and directly transferred into the slot of a blue native gel. The solubilized proteins were separated by two-dimensional Blue native/SDS PAGE as described previously by Wittig et al. (2006). The image acquisition was performed with a Typhoon Scanner, and the quantification of the Cy Dye labelled proteins resolved by the 2D Blue native PAGE was carried out using the SAMESPOTS software (Totallab, Tyne, England) based on the gel-images acquired from 3 biological replicates.

**Ruptured chloroplast assay.** The cyclic electron flow was measured in ruptured chloroplasts through an Fd-dependent plastoquinone reduction as previously described (Endo et al., 1998; Munekage et al., 2002). Briefly, the isolated chloroplasts (10 µg Chl) were osmotically ruptured in the analysis buffer (7 mM $MgCl_2$, 1 mM $MnCl_2$, 2 mM EDTA, 30 mM KCl, 0.25 mM $KH_2PO_4$, and 50 mM HEPES, pH 7.6) for 10 min prior to the analysis. One min after the initiation of the measurement, NADPH (Roche, Basel, Switzerland) was added to a final concentration of 0.25 mM. Maize ferredoxin (Sigma Aldrich, St. Louis, MO, USA) was added after an additional 30s.

**FRET-based ATP sensor and pH-dependent fluorescent probe.** The use of ATP sensor AT1.03 in plant systems was recently verified (Voon et al., 2018). A pH-dependent fluorescent probe with circularly permuted yellow fluorescent protein (cpYFP) was introduced into the matrix of the mitochondria (Nagai et al., 2001). Protonated cpYFP displays a higher absorption at 395 nm, which emits a higher signal at 528 nm if excited at 395 nm. cpYFP was shown to response specifically to the pH value (Schwarzländer et al., 2011).

**Bimolecular fluorescence complementation (BiFC).** The full-length coding sequence of the bait protein AtPAP2 was fused to the C-terminus of YFPC in the pSPYCE vector, while the coding sequences of the photosynthetic apparatus subunits were fused to the N-terminus of the pSPYNE containing YFPN (Kerppola, 2006). Both the prey and bait plasmids were transformed into the Agrobacterium tumefaciens strain GV3101, and the bacteria were infiltrated into the epidermal cell layers of tobacco leaves as previously described (Schweiger and Schwenkert, 2014). The transfected regions of the leaves were used for fluorescence detection after a 48-hour incubation in the dark under an LSM710 confocal laser scanning microscope (Zeiss). The primers that were used in this study are listed in Supplemental Table S5.

**Yeast two-hybrid assay.** A normalized yeast two-hybrid (Y2H) library was prepared from mRNAs that were isolated from eleven *Arabidopsis* tissues and was employed for the screening (Clontech Laboratories, Japan). The coding sequence of the mature bait protein AtPAP2 (25-613 a.a.), which lacks its signal peptide and C-terminal transmembrane motif, was fused to the C-terminus of the GAL4 DNA-binding domain (BD) in the pGBKT7 vector. The GAL4-based Y2H library screening was performed using the Matchmaker Gold Yeast Two-Hybrid System following the manufacturer's instructions (Clontech Laboratories, Japan). The mated culture was concentrated by centrifugation and spread onto agar plates (90 mm) containing triple dropout medium without Leu, Trp and His (TDO). Putative positive colonies (> 2 mm) were cultured on TDO liquid medium, and pGADT7 plasmids were extracted from the yeast cells for the *E. coli* transformation. The plasmids (pGADT7-prey) were re-extracted and sequenced. To identify the interactions between AtPAP2 and various components of the photosynthetic apparatus, the coding sequences of prey proteins were amplified from *Arabidopsis* leaf cDNA by Platinum$^{TM}$ *Pfx* polymerase (Thermo Scientific, USA) and fused downstream of the DNA-activating domain (AD) in the pGADT7 vector. The Y2HGold yeast cells were co-transformed with both the bait and prey protein constructs using the lithium acetate method and plated onto double dropout without Leu and Trp (DDO), TDO, and quadruple dropout without Ade, His, Leu and Trp (QDO) agar plates (Clontech Laboratories, Japan).
